# Enzyme-free nucleic acid dynamical systems

**DOI:** 10.1101/138420

**Authors:** Niranjan Srinivas, James Parkin, Georg Seelig, Erik Winfree, David Soloveichik

## Abstract

Chemistries exhibiting complex dynamics—from inorganic oscillators to gene regulatory networks—have been long known but either cannot be reprogrammed at will, or rely on the sophisticated chemistry underlying the central dogma. Can simpler molecular mechanisms, designed from scratch, exhibit the same range of behaviors? Abstract coupled chemical reactions have been proposed as a programming language for complex dynamics, along with their systematic implementation using short synthetic DNA molecules. We developed this technology for dynamical systems, identifying critical design principles and codifying them into a compiler automating the design process. Using this approach, we built an oscillator containing only DNA components, establishing that Watson-Crick base pairing interactions alone suffice for arbitrarily complex dynamics. Our results argue that autonomous molecular systems that interact with and control their chemical environment can be designed via molecular programming languages.

---

Embedded information processing circuitry provides a powerful means for creating highly functional autonomous systems. The technological capability of electro-mechanical machines has experienced revolutionary advances through embedded control. Within living organisms, embedded control is at the heart of all cellular processes, and is often seen as the distinguishing feature between living and non-living chemistries. However, in principle, non-biological chemical systems are also capable of information processing that directs molecular behaviors. Inspired by the success of systematic approaches in electrical engineering, we seek molecular building-blocks and design rules for combining them, to systematically construct non-biological autonomous molecular systems—an approach we call molecular programming.

Any candidate architecture for engineering chemical controllers must be capable of diverse dynamical behaviors. Since the discovery of well-mixed chemical oscillators (*1, 2*), synthetic reaction networks with complex temporal dynamics have been engineered based on small-molecule interactions, such as redox chemistries (*3*). The information-based chemistry underlying the central dogma of molecular biology also has been used to create various dynamical systems, such as bistable switches and oscillators in living cells (*4–7*) and in simplified cell-free systems (*8–14*) that include a limited number of enzymes. However, the range of dynamical behaviors demonstrated by synthetic systems does not yet approach the complexity and sophistication of biological circuits (*15, 16*).

Systematically engineering a wide range of dynamical behaviors would be greatly facilitated by a “programming language” for composing relatively simple molecular building-blocks into complex dynamic networks. The language of formal chemical reaction networks (CRNs)—i.e. chemical reaction equations between “formal” symbols representing species—provides a natural abstraction for specifying the diverse dynamical behaviors possible with mass-action chemical kinetics (*16–18*). Indeed, formal CRNs can be constructed to simulate arbitrary polynomial differential equations (*19, 20*), linear feedback controllers (*21*), boolean logic circuits (*22*), neural networks (*23*), distributed algorithms (*24*), and other computational models (*25*).

Dynamic DNA nanotechnology (*26*) offers an attractive molecular architecture for engineering CRNs with desired dynamic behavior. Indeed, the programmable nature of DNA-DNA interactions mediated by Watson-Crick complementarity, coupled with predictive thermodynamic models (*27, 28*) makes it possible to rationally design molecular reaction pathways (*29–33*). In particular, toehold-mediated strand displacement (*34–36*) has been exploited to engineer nanoscale tweezers (*37*), enzyme-free digital logic circuits (*38, 39*), catalytic networks (*40, 41*), and dynamically self-assembled structures (*41, 42*).

Inspired by the simple yet powerful rules governing strand displacement reactions, investigators have proposed general schemes for translating any formal CRN into a DNA implementation (*43, 44*). Confirming the validity of this approach, a general CRN-to-DNA scheme (*44*) was experimentally demonstrated for a consensus network that compared the concentration of two DNA strands and converted the “majority” into the “totality” (*45*). However, because the objective of the consensus network was a desired steady state, it remained unclear what new challenges would arise when designing dynamical behaviors.

Here, we develop a general molecular technology for engineering enzyme-free nucleic acid dynamical systems. Designing complex temporal trajectories, rather than end-point computations, places stringent requirements on kinetic design. As a challenging test case, we experimentally realized the rock-paper-scissors oscillator—a CRN that has been explored as a mathematical construct in theoretical biology (*46, 47*), ecology (*48, 49*), and molecular programming (*50*), but has not had an experimental realization in nonlinear chemistry, synthetic biology, or DNA nanotechnology. This CRN, named after the game in which “rock crushes scissors”, “scissors cut paper”, and “paper wraps rock”, consists of three autocatalytic reactions in cyclic competition. For example, from an initial state with an excess of A and only a little B and C, the reaction B + A → 2 B first autocatalytically converts most of the A to B, with an initial exponential rise in B; subsequently, the reaction C + B → 2 C quickly converts most of the B to C; and finally the reaction A + C→2 A brings the system back to the initial state.

To engineer a molecular implementation of this formal CRN, we used a CRN-to-DNA scheme based on ref. (*43*). The basic principle of this construction is to design molecular representations of each formal species that have no significant direct interactions with each other, and then for each formal reaction to design molecular machines that first detect and consume the reactants, then release the necessary products. For example, for the formal reaction U + V → X + Y, a first machine can irreversibly bind to and disable U and V only if they are both present, in which case it triggers a second machine to activate and release one copy each of X and Y. Thus, in the presence of these molecular machines, the concentrations of the molecules representing formal species follow the dynamics expected of the formal CRN, when rates are appropriately tuned.

Design of the requisite molecular machines for the rock-paper-scissor oscillator followed a systematic hierarchical strategy. In the first stage, the machines were designed according to a so-called “domain-level” model that captures the generic biophysics of DNA association, dissociation, and branch migration. In the second stage, specific nucleotide sequences were chosen with the goal that they approximate the behavior of the domain-level model as closely as possible. Synthesized, tested, and debugged in the laboratory, we verified the functionality of each autocatalytic module, demonstrated the full oscillator, and quantitatively characterized the system. In doing so, we identified several critical design principles, both at the domain level and at the sequence level, and acquired improved understanding of molecular non-idealities, including means of mitigating and compensating for imperfect execution of desired reactions and spurious “leak” reactions. Our mechanistic model with measured kinetic parameters provides a proof by synthesis that designed molecular interactions are sufficient for programming mass-action kinetics. Lastly, we implemented a CRN-to-DNA “compiler” that incorporates our design principles: given a formal CRN, it automates the design process to provide candidate DNA sequences for implementing the desired dynamical behavior. Every experiment performed with DNA sequences designed by our automated compiler led to oscillatory dynamics on the first try. Our successful demonstration of a general technology for building chemical dynamical systems may enable systematic engineering of other sophisticated dynamical behaviors, which may act as controllers for the ever-expanding range of molecular structures, machines, and devices developed in DNA nanotechnology.

## CRN to DNA implementation scheme

Given a dynamical behavior as specified by a formal CRN program, we aim to systematically design a DNA-based implementation that approximates the specified behavior in a test tube (Fig. 1A). We represent each formal species in the CRN program by a set of single-stranded DNA species (“signal strands”). These signals are designed not to interact directly with each other. Instead, each reaction in the formal CRN is mediated by additional DNA species (“fuels”) that provide both logical machinery, free energy, and material for the desired reaction to occur. This strategy is modular because implementing one more formal reaction only requires adding the corresponding set of fuel species. Signal strands correspond to formal species in the following sense. Signal strands can be either active, or inactivated when bound in a larger complex. The temporal evolution of the concentrations of active signal strands is expected to approximate the kinetics of the corresponding formal species. The approximation holds as long as the fuel species remain sufficiently in excess because of a mathematical argument based on singular perturbation theory (*43*). Our experiments were done in a one-pot batch-reactor, without any flow of matter or energy; therefore, the dynamics of our test-tube realizations were expected to deviate from the specified dynamics once a substantial fraction of fuel species was consumed.

**Fig. 1.**
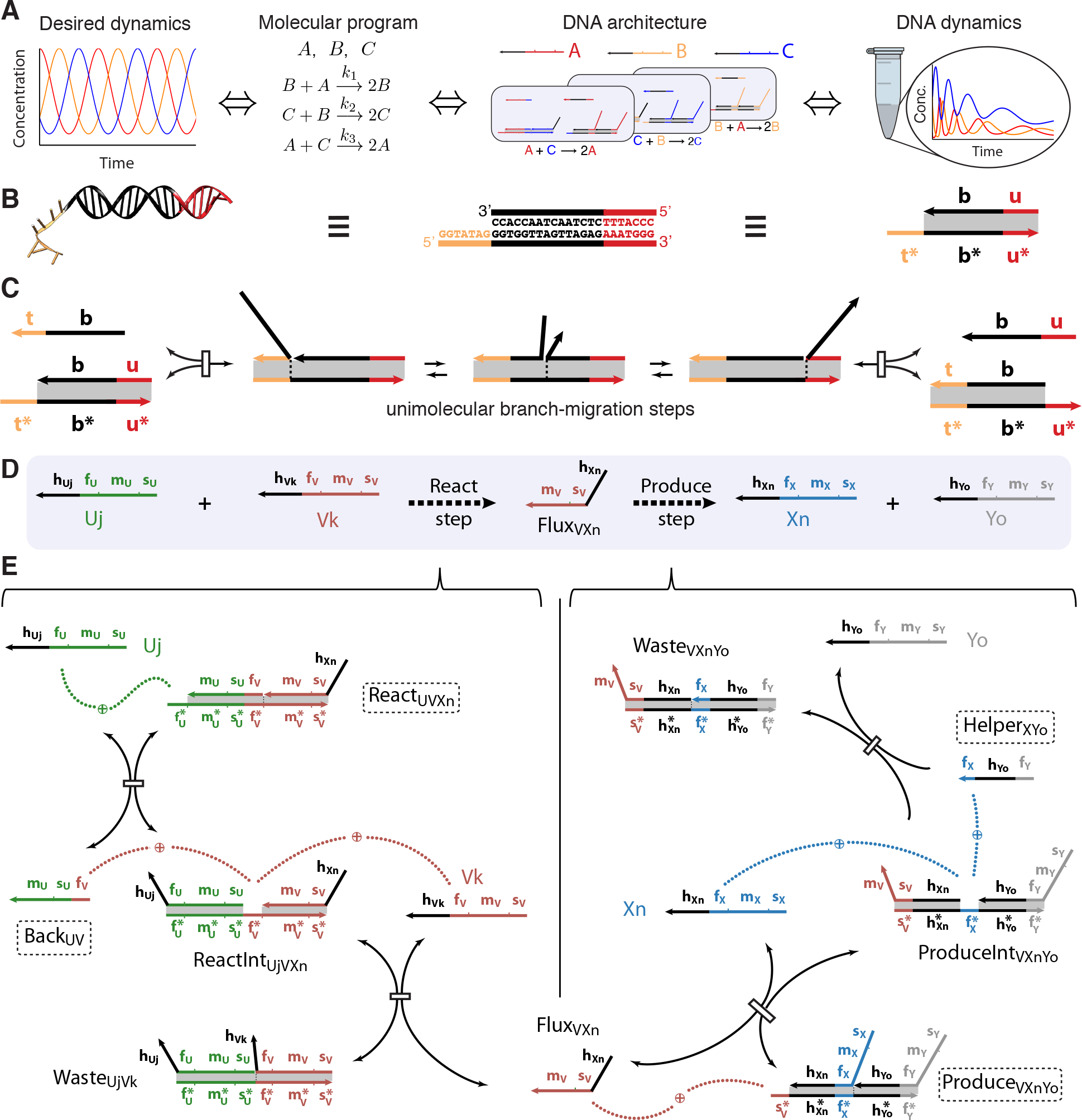
A systematic pipeline for engineering dynamical systems with DNA strand displacement cascades. (**A**)Desired dynamics (oscillation) are specified by a molecular program (three-reaction “rock-paper-scissors” CRN over formal species A, B, C, with rate constants *k*_1_, *…, k*_3_), and is instantiated using our DNA architecture. A set of DNA strands called signal strands corresponds to the formal species in the following sense: the temporal evolution of the amount of signal strands A that is free (active) approximates the kinetics of the corresponding formal species A (equivalently for B, C). The implemented temporal kinetics (schematically illustrated under “DNA dynamics”) are driven by certain meta-stable DNA complexes and strands (collectively called fuels), and dynamics is expected to diminish as the fuels get consumed. (**B**) Domain-level abstraction of a multi-strand DNA complex. Domains are contiguous sequences that act as a unit; names t, b, u are shorthand for those sequences. Arrows indicate 3′ ends; ∗ denotes Watson-Crick complementarity. (**C**) Building-block strand displacement interaction. Open toehold domains t and t^*∗*^ bind, followed by displacement of domain b of strand b–u by strand t–b, followed by dissociation of short domain u. (**D**) High-level implementation of the general bimolecular reaction U + V → X + Y. In the react step signal strands U and V are bound (inactivated), and in the produce step signal strands X and Y are released (activated). The react and produce steps are mediated by fuel species (shown in E), and are coupled through an intermediate Flux strand. (**E**) Strand displacement implementation of the react and produce steps. Fuel species are labeled by dashed boxes. Dotted arcs illustrate toehold binding and dissociation interactions. Subscripts on signal strands in parts D and E correspond to different history domains h, which are used to hold signal species inactive in a Produce complex, but have no role once the signal strand is released.

We now describe the domain-level abstraction that facilitates our hierarchical design, explain the strand displacement interaction that serves as a building-block, and finally illustrate the strand displacement steps that implement formal CRN reactions.

As part of our hierarchical design strategy, we use logical constructs called “domains”, which are contiguous base positions that are intended to act as a unit (Fig. 1B). We design interactions entirely at the domain level, and later generate concrete sequences that are expected to satisfy the domain level specification. The basic strand displacement interaction (*34–36*) is illustrated in Fig. 1C, where two single strands of DNA compete for binding with their partial complement. Each competing strand shares a long (13-25 nt) “branch-migration” domain that is bound strongly enough that spontaneous dissociation does not occur. The reactions are mediated by short (5-7 nt) single-stranded domains called “toeholds” that fleetingly co-localize the competing strands, thereby facilitating the in tramolecular exchange of base pairs (branch migration). Eventually, one of the competing strands dissociates. Specificity is achieved by the choice of DNA sequence; many orthogonal strand displacement interactions can occur simultaneously in the same solution as long as sequence overlap is minimized.

As illustrated for the general bimolecular reaction U + V→ X + Y in Fig. 1D, the implementation of a formal reaction is conceptually divided into two steps: (1) the “react step” recognizes and consumes the reactants U and V as input, and (2) the “produce step” releases the products X and Y as output. These steps are linked by a Flux strand, released in the first step, that triggers the second step. Each step occurs as a sequence of strand displacement interactions driven by the fuel species described below.

A formal species (e.g. U) is represented by a set of signal strands (Ui, Uj, *…*) which are identified by three common domains: the first toehold (f_U_), the branch migration domain (m_U_) and the second toehold (s_U_). All desired strand displacement interactions involving a given formal species (U) occur with these three domains. Additional “history” domains (h_Ui_, h_Uj_, *…*) which can vary between strands corresponding to the same formal species (Ui, Uj, *…*) are intended to be inert; they merely facilitate the formation of the desired fuel structures (specifically when the Produce complexes are annealed, Fig. S1).

The mechanism of the react step shown in Fig. 1E (left) begins with the React complex (React_VUXn_) reversibly binding and inactivating a signal strand representing the first formal reactant (U), releasing the Backward strand (Back_UV_). Because the React complex and Backward strand are both in excess (both are fuels), the resulting intermediate complex (ReactInt_VUXn_) is expected to approach a pseudoequilibrium proportional to the concentration of U. Thus, ReactInt_VUXn_ interacts with the second input, V, at a rate proportional to the product of their concentrations, in accordance with standard mass-action chemical kinetics for bimolecular reactions. This reaction irreversibly releases the Flux strand (Flux_VUn_).

In the subsequent produce step shown in Fig. 1E (right), the Flux strand initiates a pathway that releases products X and Y, which are initially bound to the Produce complex (Produce_VXnYo_) by their history domains and first toeholds. After the Flux strand releases the first output, a toehold is exposed that allows the Helper strand (Helper_XYo_) to irreversibly displace the second output. Because each forward reaction is driven by a fuel species that is in high concentration, the produce step reactions are fast relative to the rate-limiting react step, which therefore determines the overall pathway kinetics.

Our CRN-to-DNA scheme is fully general: it can be used to construct reactions with repeated reactants or products (as we demonstrated with autocatalysis) or a different number of reactants and products (e.g. unimolecular reactions can be obtained by using a single-stranded fuel species as one of the reactants). Our naming scheme is both precise and general—the name and the molecule fully determine each other (Fig. S2), which facilitates automated and systematic analysis (Note S6).

## Non-idealities in a single-reaction CRN

To understand the challenges in using our general CRN-to-DNA scheme for engineering dynamical behaviors, we used the autocatalytic single-reaction CRN C + B→ 2 C as a test case (Fig. 2A). We obtained the domain-level specification of the fuels for driving the autocatalytic reaction by starting with the general prescription shown Fig. 1E, and replacing the domains of U, V, X, and Y by those of C, B, C, and C respectively (Figs. 2B, S3). Since the kinetics of exponential amplification is sensitive to both the initial concentration of the autocatalyst and the effective rate constant of the reaction, this system provided a stringent test of our ability to control dynamics.

**Fig. 2.**
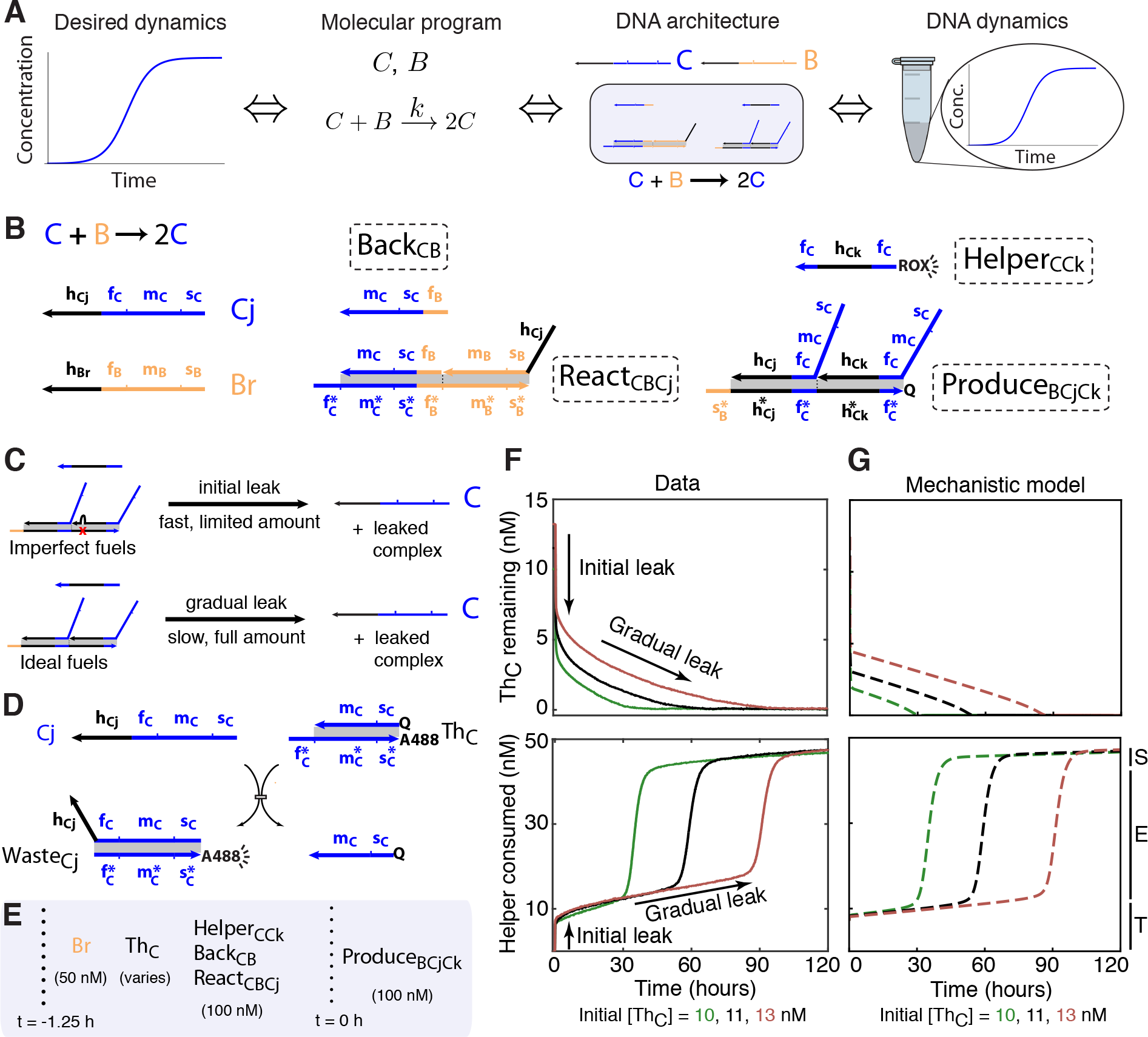
The implementation of a single reaction autocatalytic CRN, and counteracting non-idealities. (**A**)Engineering schematic using our systematic pipeline. (**B**) DNA species which needed to be synthesized. The domain level design followed from the prescription of Fig. 1D, E (fuel species are indicated by dashed boxes). Two sources of leak. A limited amount of imperfect fuel molecules, such as those with DNA synthesis errors, release signal strands and waste products through fast spurious pathways (initial leak). Ideal fuel molecules release similar products through slow gradual leak. (**D**) Counteracting initial leak. A Threshold complex (Th_C_) consumes a stoichiometric amount of leaked autocatalyst C. (**E**) Experimental setup. Vertical dotted lines separate initial contents of the test tube and timed additions. Addition of Produce complexes kickstarts release of autocatalyst through initial and gradual leak. (**F**) Experimental data showing the concentration of Th_C_ (top) and the amount of Helper_CCk_ consumed (bottom) for three independent samples with differing initial amounts of Th_C_. Reaction progress can be logically split into three phases: The threshold consumption phase (T) is dominated by leak consumption; the exponential phase (E) is dominated by the intended autocatalytic reaction, and initiates when threshold is exhausted; and the saturation phase (S) refers to the completion of the intended reaction. The progress of the reaction was monitored via fluorophores (ROX, A488) on the Helper and Threshold species shown in B and D, which emit measured signal when separate from the quencher (Q). (Notes S4.1, S8.) (**G**) Mechanistic model semi-quantitatively captures the dynamics of the DNA implementation (Note S5).

Experiments on the single-reaction CRN as well as the full oscillator were performed similarly (see S8 for details). DNA strands were chemically synthesized, and fuel complexes were individually annealed and gel-purified to ensure the correct stoichiometry of the constituent strands. To observe reaction progress, we incorporated fluorophores and quenchers into certain molecules (e.g. Helper and Produce fuels in Fig. 2B). The monitored strand displacement interactions resulted in the co-localization of the fluorophore and quencher, which suppressed fluorescence (Notes S4.1). Spectrofluorimetry experiments were performed in TE buffer (Tris/EDTA, pH 8.0), with 0.5 M NaCl at 25^*°*^C.

The success of an experimental realization is determined by how well the system behaves in accordance with the domain-level model. DNA strand displacement systems suffer from three main classes of molecular non-idealities: output strands can be released when they should not be (“leak”), input strands can be consumed without producing output (“sub-stoichiometric yield”), and reactions can proceed at the wrong rate.

We observed both a limited amount of fast “initial leak”, as well as a slower “gradual leak” that continued throughout the duration of the experiment (Figs. 2C, F; S16). Initial leak is thought to arise primarily from a fraction of imperfectly prepared fuel molecules—e.g. due to synthesis errors (truncations or deletions) in individual strands, or improperly folded multi-stranded complexes—that, when initially mixed together, can readily interact and release their outputs. In contrast, gradual leak cannot be avoided even with perfectly synthesized and folded molecules, because it arises from the inherent biophysics of strand displacement (*36*). Mechanistically, these reactions could initiate from invasion at the end of a helix (blunt-end) or a coaxial junction even in the absence of a toehold (*51*) (Fig. S6), or could be facilitated by spurious remote toeholds (*52*) (Fig. S19, S20).

Substochiometric yield can arise as a consequence of leak pathways: both initial and gradual leak can result in reactive complexes that can consume inputs without releasing outputs, because the outputs have already been released (Fig. S7). Substoichiometric yield can also result from other synthesis errors—e.g. truncations on toehold regions of output strands could render them nonfunctional for triggering downstream reactions (Fig. S8).

Desired reactions can take place with markedly different kinetics because of sequence differences, even when the domain-level descriptions are identical. This is partly due to the exponential dependence of strand displacement rate constants on toehold length and binding energy (*34, 35*), but can also be affected by undesired secondary structure within signal strands as well as fleeting binding between toeholds and unrelated single-stranded portions of unrelated molecules, both of which can occlude the toehold and inhibit the desired reaction.

Both types of leak were clearly evident in our experimental implementation of the single-reaction autocatalytic CRN that converted an initial reservoir of B into C, with the fuels in excess. To prevent the immediate onset of exponential amplification from the initial leak of the autocatalyst, C, we introduced a Threshold complex (Th_C_) designed to quickly consume C (Fig. 2D). At an initial Threshold concentration greater than the initial leak of C, exponential amplification was delayed until further gradual leak of C eventually exhausted the Threshold. At that point, further gradual leak triggered exponential amplification of C by the implemented reaction C + B → 2 C, until the provided quantity of B was fully consumed. Monitoring the progress of the reaction, with fluorophores positioned on the Threshold and Helper species, showed the expected proportional delays for three different initial amounts of Threshold (Fig. 2F). Qualitatively similar delayed amplification was also seen in two other single-reaction autocatalytic CRNs, A + C → 2 A and B + A → 2 B (Figs. S4, S5, S11, S12). The three modules had different initial and gradual leak rates (Fig. S14, Table S3, and Note S7.2), resulting in a roughly 10-fold variation in the delay times (Fig. S13).

Evidence for substoichiometric yield and non-ideal reaction rates can also be seen in Figs. 2F, S13 (e.g. the autocatalytic phase did not consume the full complement of 50 nM of Helper, and the amplification rates were not equal for the different delays), but the clearest evidence came from measurements of individual steps of the reaction pathways. For analogous reactions, rate constants were all within a factor of 20 of each other, and mostly within a factor of 3 (Tables S1, S2). Substoichiometric yields on the order of 20% lower than ideal were observed (Fig. S22).

The essential features of the autocatalytic dynamics were captured by a quantitative mechanistic model at the level of individual strand displacement reactions (Fig 2G). For each of the three autocatalytic CRNs, all 7 rate constants for reactions shown in Figs. 1E and Fig. 2D were measured separately (Tables S1 and S2). In addition to the 21 independently measured rate constants for the desired reaction pathways, the model partially accounted for observed non-idealities (Note S5). The Produce-Helper leak (Fig. S6) was determined to be the dominant pathway and included in the model; the 3 rate constants for gradual leak were inferred from the 3 autocatalytic module experiments (Fig. S14, Table S3, and Note S7.2), whereas the amount of initial leak was assumed to be the same for all modules, and was specified by a single parameter. Sub-stoichiometric yield resulted from interactions with the products of leak reactions (Fig. S7) as well as an assumption that a fraction of output strands (Flux and Signal molecules) were non-functional (Fig. S8, the same fraction in all cases). Finally, brief toehold binding in positions where strand displacement cannot occur (toehold occlusion) can cause a concentration-dependent decrease in strand displacement rates (*39*). To account for toehold occlusion we introduced a single parameter for the strength of toehold binding in all unproductive contexts (Fig. S9).

The three empirical parameters (for initial leak, non-functional output, and toehold occlusion) were fit to the full oscillator data of Fig. 4B, including an additional fitting parameter for each initial signal concentration to account for imperfect pipetting and uncertainties in the initial leak. Removing any of these non-idealities from the mechanistic model prevented adequate explanation of the data. We used the same three empirical parameter values when modeling the three single-reaction CRNs, using only three newly-fit parameters for the Threshold concentrations, to account again for uncertainties in the initial leak. We interpret the success of the mechanistic model to mean that we captured the dominant effects in our system, and we expect that similar models will have considerable predictive power for future strand displacement systems.

## Principles for robust molecular design

The previously discussed understanding of non-idealities in strand displacement cascades was refined over the course of four iterations of sequence design for the three autocatalytic modules, in parallel with the development of sequence design principles and experimental methods that helped minimize the non-idealities (Notes S3, S4). The performance of Design 4, presented above, is an improvement over each previous design (Note S4).

Initial leak was reduced by several design choices and experimental methods. First, the baseline sequence design criterion was sequence symmetry minimization (*29*), which unlike purely thermodynamic approaches (*31*) is expected to help the folding process avoid being kinetically trapped in malformed conformations (*30*). Second, fuel complexes were prepared by annealing HPLC-purified oligonucleotides, followed by PAGE gel purification to minimize undesired multimers and excess single-strands (Note S8). Third, because the orientation of bases on the DNA backbone (5^*1*^-3^*1*^) is known to affect the distribution of synthesis errors (*53*), we achieved a three-fold reduction in initial leak by using the backbone orientation in which the toehold occurs on the 5^′^ end (Note S3.5). Lastly, Threshold complexes were used to tune the initial conditions by removing leaked signal strands from solution.

Since gradual leaks primarily arise from strand displacement through invasion at frayed blunt ends and coaxial junctions, we used 2-nt clamps at the end of React and Produce complexes and closed helices and coaxial junctions with strong (C/G) base pairs (Fig. S15). Further, we minimized spurious remote-toehold strand displacement (*52*) by avoiding even relatively weak complementarity at overhangs near coaxial junctions (Fig. 3). These strategies reduced gradual leaks as much as 15-fold relative to earlier designs (Table S5, Fig. S16, Fig. S18).

**Fig. 3.**
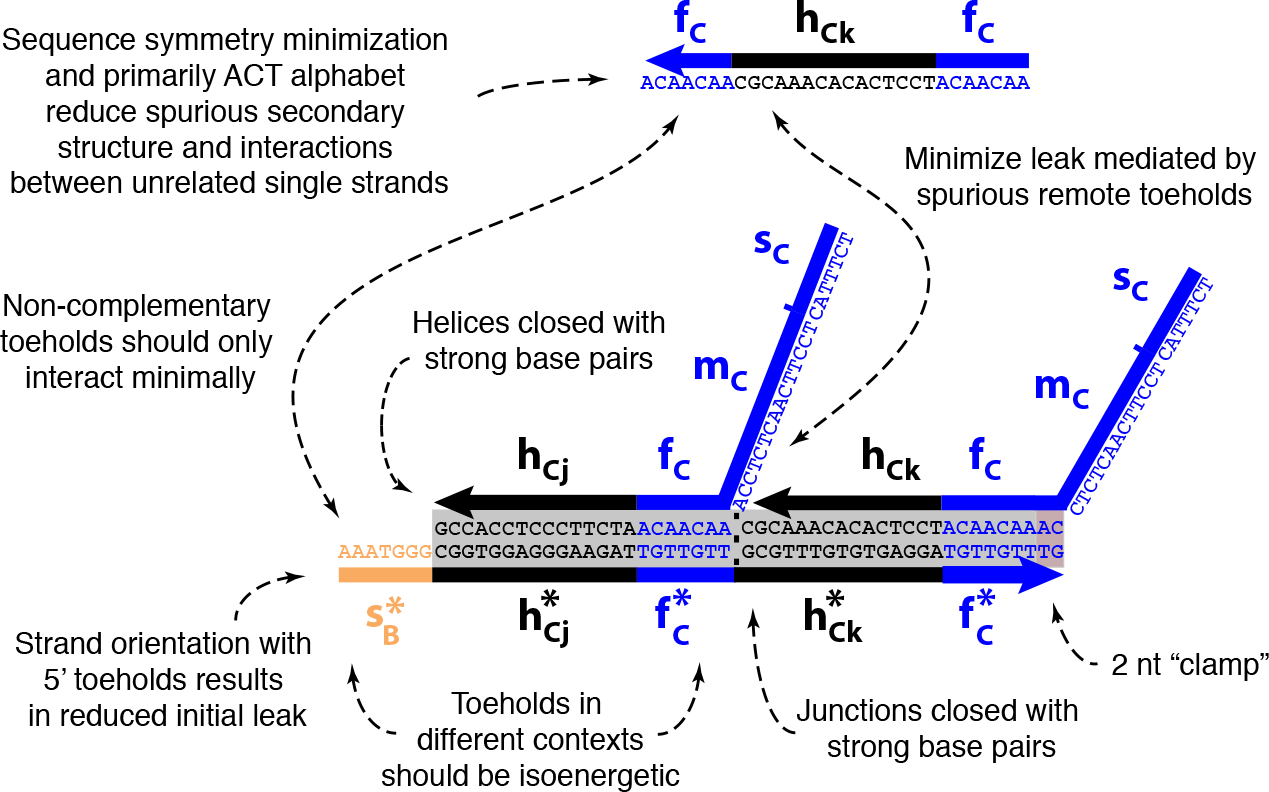
Sequence design principles illustrated with a Produce complex (Produce_BCjCk_).

Three key design strategies were used to minimize undesired variability in rates. First, signal strands contained at most one guanine, reducing the propensity for promiscuous intramolecular base-pairing (*35, 39, 40*); in particular, we ensured that toeholds and the first 4 bases of branch migration domains, which are crucial for initiating strand displacement, were free of unintended secondary structure (Fig. S21, Note S3.6) (*35, 36*). Secondly, toe-holds were designed to be isoenergetic according to nearest-neighbor parameters (*27*) augmented with terms for coaxial stacking and protruding tails at nicks (Note S6.2) (*35, 36*). Because the same toeholds occurred in different contexts—with different flanking structures and thus different energetics—we truncated 1 or 2 nucleotides in some cases to help equilibrate binding energy (specifically in the reversible toehold exchange step in the React complex pathway, which is expected to be rate determining, Fig. S22). Finally, toeholds were simultaneously designed to be as orthogonal as possible, and branch migration domain sequences were designed to be orthogonal to toeholds, in order to mitigate toehold-occlusion and the concomitant decrease in strand displacement rates (Note S5.3, Table S4) (*39*).

To combat substoichiometric yield, we designed a modified Helper strand that, in addition to displacing the second output in the produce step, also displaces the original Flux strand that initiated the produce step, effectively enabling catalytic action by the Flux strand (Fig. S10). This “catalytic Helper” permits the Flux strand to release more outputs by initiating displacement with another Produce molecule, thereby increasing effective reaction stoichiometry. By tuning the relative concentration of the catalytic Helper strands, we adjusted reaction stoichiometry—much as potentiometers were used to tune the performance of early electrical circuits.

The design principles used to curtail non-idealities involved an unusual combination of thermodynamic, kinetic, and ad-hoc criteria, which were not compatible with straightforward application of state-of-the-art sequence design tools (*28*). Therefore, we formulated custom heuristic measures for comparing candidate sequence designs and implemented them as a collection of scripts that called NUPACK (*28*), Pepper (*54*), Sticky-Design (*54, 55*), and SpuriousSSM (*54*), in order to perform sequence design and analysis (Note S3). Because the three autocatalytic modules were intended to work together as an oscillator, as described in the next section, they were designed together as a single system.

## A DNA strand displacement oscillator

The strand displacement oscillator (which we call the Displacillator, Fig. 4A) realizes the rock-paper-scissors CRN (*46–50*). The orbit of this neutral cycle oscillator is determined by conservation laws for two quantities: [A] + [B] + [C] and [A]^*k*_2_*/k*_0_^ [B]^*k*_3_*/k*_0_^ [C]^*k*_1_/*k*_0_^, where [A], [B], [C] are the concentrations of the three formal species, *k_i_* the rate constants of the three formal reactions, and *k_i_/k*_0_ are unitless rate constants (*48*). Unlike most limit cycle oscillators, the rock-paper-scissors CRN oscillates for any choice of reaction rate constants and (non-steady-state) initial signal concentrations, which makes it especially suitable for implementation as a dynamical strand displacement cascade. The number of oscillation periods expected before fuels are exhausted decreases with the amplitude of the oscillation. Thus, to reduce the amplitude resulting from initial leak, we added 10 nM of each Threshold complex to consume all the spuriously released signal strands, yielding a metastable mixture of fuels with at most 2.5 nM of each Threshold complex left over (Fig. S23). To initiate the reactions underlying the Displacillator, we then added signal strands in excess of the residual Thresholds to target a suitable initial state of the oscillator (Fig. 4B). The concentration of catalytic Helpers, added to compensate for substoichiometric yield, was empirically tuned to 25% of total Helper concentration. (Fig. S23 illustrates experimental setup.)

**Fig. 4.**
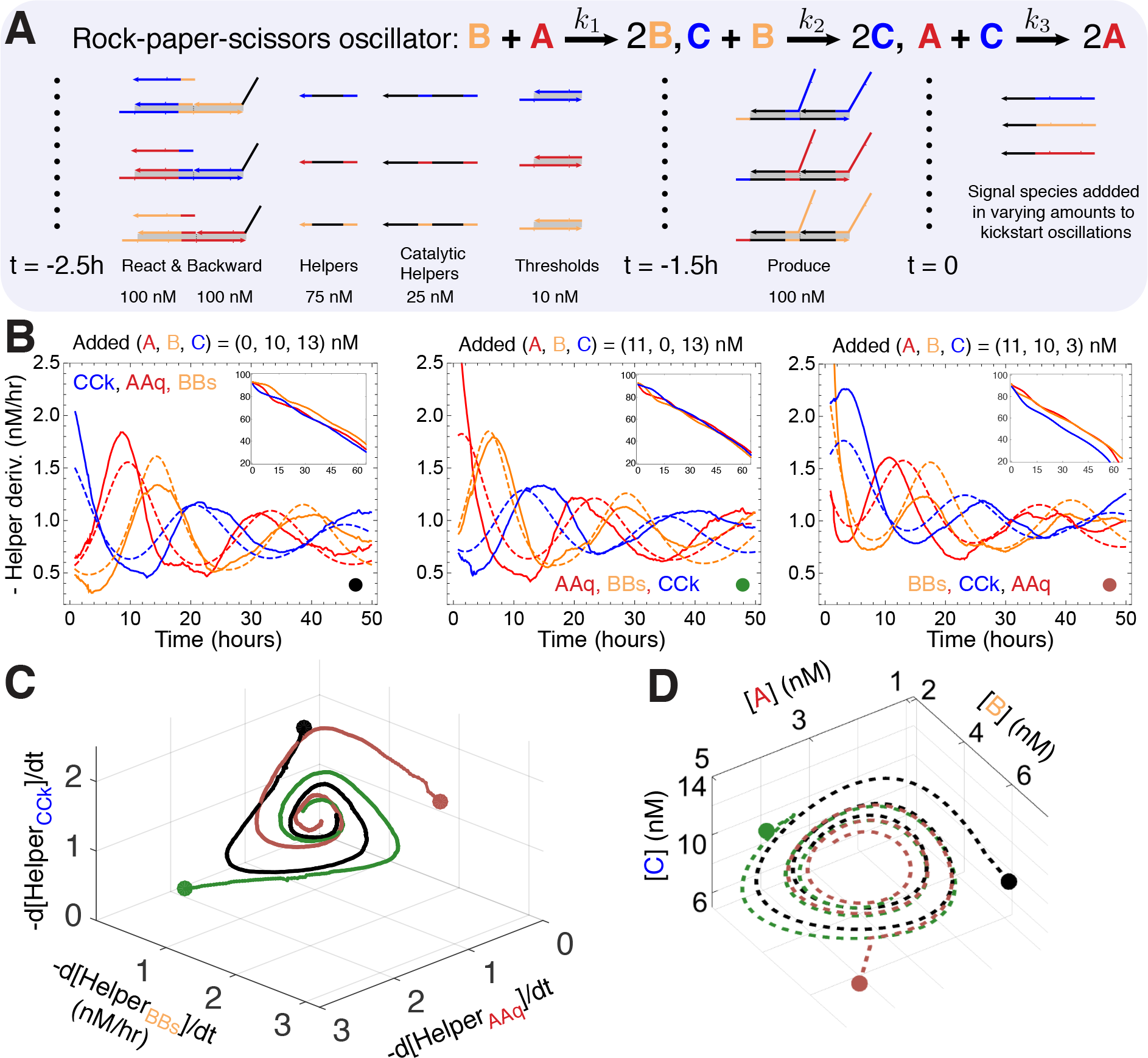
Experimental realization of a strand displacement oscillator (Displacillator). (**A**) The rock-paper-scissors CRN, and the schematic of the DNA species which needed to be synthesized following the prescription of Fig. 1D, E. Vertical dotted lines separate initial contents of the test tube and timed additions. (**B**) Experimental data (solid lines) and mechanistic model fits (dashed lines) show time derivatives of the consumption of the three Helper strands under three different initial conditions. Insets display measured Helper concentrations (in nM). (**C**) Phase plot of the experimental data shown in B. Thick dots indicate initial conditions. (**D**) Phase plot of the concentrations of the signal strands extrapolated from the mechanistic model.

Observing system dynamics by directly measuring free signal strand concentrations may consume or temporarily sequester a fraction of these strands. Instead, to avoid interfering with the dynamics, we measured the net progress of each reaction by monitoring the quenching of fluorophore-labeled Helper strands. Oscillatory dynamics were observed in the instantaneous consumption rates of the Helper strands (Figs. 4B, C; S24), until the fuel species were depleted. The order in which the reaction rates reached peaks and minima was consistent with the ideal rock-paper-scissors dynamics, for each of the 3 initial concentrations of signal species.

The mechanistic model (see above and Note S5) demonstrates that the emergent dynamics of the reaction mixture can be quantitatively explained by the individual strand displacement interactions that we designed, and the non-idealities that we understand. The mechanistic model accounted for most of the measured Helper consumption dynamics, including the eventual decrease in rate due to fuel depletion (Fig. 4B). The model further allowed us to extrapolate signal (A, B, C) concentrations that were not directly accessible to measurement (Fig. 4D). Although we observed oscillations in reaction rates directly by measuring Helper fluorescence, the extrapolated signals allowed us to tie A, B, C signal dynamics back to the ideal rock-paper-scissors CRN (Note S5.5, Fig. S26). This agreement with the design specification—the formal CRN— confirmed that the Displacillator oscillated for the reasons that we intended.

## A CRN-to-DNA compiler

In principle, our design strategies could be used to construct an automated pipeline for implementing any formal CRN. To do so, we integrated the sequence design tools and principles into an end-to-end compiler, called Piperine, that accepts an arbitrary formal CRN as input and produces candidate sequences for experimental implementation. Piperine’s sequence design pipeline proceeds in several stages: first, the formal CRN is translated into a set of requests for the necessary fuel molecules in the Pepper DNA design-specification language (*54*); second, Pepper uses templates for each type of fuel strand or fuel complex to deduce the full set of strands and base-pairing constraints; third, toeholds are designed to be orthogonal and isoenergetic using StickyDesign (*54, 55*); fourth, the toehold sequences and base-pairing constraints are sent to SpuriousSSM (*54*), which uses sequence symmetry minimization to obtain sequences for the long domains; fifth, proposed sequence sets are scored using heuristic criteria that make use of NUPACK (*28*) to evaluate key secondary structure and spurious binding interactions; finally, multiple independent sequence designs are compared according to these criteria, and the sequence set that scores best across the board is recommended (Note S6.3).

In order to test the compiler, we used it to design, from scratch, another instance of the Displacillator with completely independent sequences. Every experiment performed with these independent sequences, from each of the three autocatalytic modules to the full oscillator, worked the very first time (Fig. S35, Note S7), leading to an independent DNA-only oscillator in just 4 weeks from sequence design to experimental demonstration. When applied to other CRNs of comparable size, it is reasonable to expect that Piperine will produce sequences that perform well for implementing other dynamical systems. More generally, it would be straightforward to augment Piperine to compile CRNs using other translation schemes (*43–45, 50, 56, 57*). Indeed, the core sequence design principles used here heuristically (depicted in Fig. 3) could serve as a valuable reference point for the development of rigorous sequence design methods, incorporating both thermodynamic and kinetics constraints, for an even wider range of strand displacement cascades (*33*).

## Conclusions

The development of programmable molecular technologies will require systematic architectures and automated design software. Our demonstration of a chemical oscillator using just DNA strand displacement cascades prototypes such a general technology for chemical dynamical systems. We expect that our molecular design principles and experimental methods can be used to implement molecular programs with diverse temporal trajectories, utilizing no more than the principles of Watson-Crick base pairing. The well-understood molecular mechanisms underlying DNA strand displacement (*34–36*) permit detailed mechanistic design of reaction pathways, which in turn enables quantitative modeling at the level of individual strand displacement reactions.

Dynamical systems, including oscillators, instantiated in biochemistry and programmed by the choice of DNA sequence have at least a 20-year history (*8, 9, 59, 60*). In contrast to prior work, a motivating principle for our simple DNA architecture was to design all molecular components from scratch, using a detailed mechanistic understanding of how and why they work. In particular, we avoided “black box” molecular components that have not been rationally designed, such as enzymes. One consequence is that, compared to other in-vitro biochemical oscillators that do make use of enzymes, the Displacillator has lower overall complexity when the complexity of the enzymes is taken into account (Table 1). For example, the “genelet” architecture (*9, 10, 61*) simplifies genetic regulatory networks (GRNs) by avoiding protein synthesis and using RNA to directly regulate transcription from short DNA templates; it relies on two essential enzymes, an RNA polymerase and a ribonucle-ase. The PEN toolbox architecture (*59*) goes further by also eliminating RNA altogether, using just a DNA polymerase, an exonuclease, and a nickase. Finally, cell-free transcription-translation (TX-TL) architectures (*60*) are sufficient for implementing many GRNs without the full complexity of living cells; whether derived from cell extract or reconstituted from purified components (*62*), over 100 essential components are involved (polymerases, ribosomes, tRNA, tRNA synthetases, amino acids, NTPs, etc). In each of these architectures, many different circuits can be implemented by introducing suitably designed DNA molecules. For the reference oscillators, we use the number of designed nucleotides as a simple metric for design size, and the total number of base-pairs of DNA that code for enzymes as a proxy for the extent of black-box genetic information. Although the Displacillator has the lowest overall design complexity compared with the other oscillators in Table 1, its relatively poor performance highlights the remaining challenges for fully rationally designed biochemical dynamical systems.

**Table 1.** Comparison to other recent synthetic cell-free biochemical oscillators. ^*†*^ T7 RNA polymerase, E. coli Ribonuclease H, and pyrophosphatase; ^*‡*^ Bst DNA polymerase, RecJf exonuclease, and Nt.BstNBI nickase.

| Oscillator | Design size | Black-box size | Number of Cycles | Period |
| --- | --- | --- | --- | --- |
| DSD: Displacillator | 1386 nt | 0 | 3 (batch) | ~ 20 h |
| Genelets: Design I (10) | 469 nt | ~ 4000 bp<sup>†</sup> | 6 (batch) | ~ 3 h |
| PEN: Oligolator (11) | 71 nt | ~ 7700 bp<sup>‡</sup> | 30 (batch) | ~ 1.7 h |
| GRNs: Repressillator (5, 58) | 6664 bp | > 100 components | ≫ 9 (microfluidics) | ~ 3 h |

There are currently many proposals, some partially demonstrated, for implementing CRNs with DNA (*43–45, 50, 56, 57*). Each scheme makes different choices regarding the representation of signals and the implementation of desired reactions, resulting in different molecule sizes, number of additional mediating species, lengths of reaction pathways, sequence design constraints, and potential for leak reactions. It is not clear how to best compare these schemes in terms of their potential for engineering arbitrary dynamical behaviors in the test tube. Improved understanding of the biophysics of initial and gradual leak pathways, and of the sequence-dependence of kinetics for fundamental DNA mechanisms such as hybridization, branch migration, fraying, and dissociation (*36, 63, 64*), should allow molecular systems to be designed with more accurate control over kinetics and with less leak. Indeed, certain CRN-to-DNA schemes may have orders-of-magnitude lower leak (*65*), raising the prospect that higher concentrations and thus faster kinetics could be achieved reliably. Providing a continuous “power supply” by replenishing fuel species and removing waste molecules (as in a continuous-flow stirred reactor (*66*)) could enable faithful dynamics on longer time scales, such as those required for controlling self-assembly or chemical reactors.

Enabling the reliable and routine use of enzyme-free nucleic acid dynamical systems as embedded chemical controllers will require integrating nucleic acid subsystems with a broad range of other chemical processes. Strand displacement cascades already have enhanced potential for modular integration with the ever-expanding range of molecular structures, machines, and devices developed in DNA nanotechnology (*67, 68*). Further-more, nucleic acids—both DNA and RNA—are well known for their ability to bind to and sense small molecules (*69, 70*), thus providing direct mechanisms to “read” the chemical environment. Nucleic acid nanotechnology has also been applied to control chemical synthesis (*71–73*); to control the arrangement (and rearrangement) of metal nanoparticles, quantum dots, carbon nanotubes, proteins, and other molecules (*74–78*); and to control the activity of enzymes and protein motors (*79–81*). Much as genetic regulatory networks and other biochemical feedback networks control chemical and molecular functions within biological cells, it is conceivable that nucleic acid dynamical systems could serve as the information processing and control networks within complex synthetic organelles or artificial cells (*82*) that sense, compute, and respond to their chemical and molecular environment.

## Acknowledgments

The authors thank L. Qian for help with initial experiments and technical training, C. G. Evans for supporting the development of Piperine, A. Phillips for assistance with Visual DSD, and C. Geary, C. T. Martin, N. A. Pierce, P. W. K. Rothemund, S. L. Sparvath, B. Wolfe, F. Dannenberg, and D. Y. Zhang for helpful discussions. This work was supported by NSF Grants 0728703, 0829805, 0832824, 1317694, 1117143, ONR award N00014-16-1-2139, the Gordon and Betty Moore Foundation’s Programmable Molecular Technology Initiative, and NIGMS Systems Biology Center grant P50 GM081879. Data presented in the paper are archived in the Supporting Online Material.

## Supplementary materials

Figs. S1-S74, Tables S1-S17, and References 83-92 are available at Science online http://science.sciencemag.org.

